# Circulating Levels of Free 25(OH)D Increase at the Onset of Rheumatoid Arthritis

**DOI:** 10.1101/675124

**Authors:** Vidyanand Anaparti, Xiaobo Meng, Hemsekhar Mahadevappa, Irene Smolik, Neeloffer Mookherjee, Hani El-Gabalawy

## Abstract

**Objective:** Epidemiological studies suggest vitamin D deficiency as a potential risk factor for rheumatoid arthritis (RA) development, a chronic autoimmune disorder highly prevalent in indigenous North American (INA) population. We therefore profiled the circulating levels of 25-hydroxyvitaminD [25(OH)D], an active metabolite of vitamin D, in a cohort of at-risk first-degree relatives (FDR) of INA RA patients, a subset of whom subsequently developed RA (progressors).

**Methods:** 2007 onward, serum samples from INA RA patients and FDR were collected at the time of a structured baseline visit and stored at −20°C. Anti-citrullinated protein antibodies (ACPA), 25(OH)D, hs-CRP, vitamin-D binding protein (VDBP) levels were determined using ELISA and rheumatoid factor (RF) seropositivity was determined by nephelometry.

**Results:** We demonstrate that 25 (OH) D concentrations were lower in winter than summer (*P*=0.0538), and that serum 25(OH)D levels were higher in samples collected and stored after 2013 (*P*<0.0001). Analysis of samples obtained after 2013 demonstrated that 37.6% of study participants were 25(OH)D insufficient (<75nmol/L). Also, seropositive RA patients and FDR had lower 25(OH)D levels compared to ACPA-/FDR (*P*<0.05, *P*<0.01 respectively). Linear regression analysis showed 25(OH)D insufficiency was inversely associated with presence of RA autoantibodies. Longitudinal samples from 14 progressors demonstrated a consistent increase in 25(OH)D levels at the time they exhibited clinically detectable joint inflammation, without any significant change in VDBP levels.

**Conclusion:** We demonstrate that 25(OH)D levels in serum increased at RA onset in progressors. The potential role that vitamin D metabolites and their downstream effects play in RA transition requires further investigation.

## INTRODUCTION

RA is a systemic autoimmune inflammatory disorder that disproportionately affects indigenous North American (INA) population (1). The reasons for this are incompletely understood and appear to relate to complex gene-environment interactions that initiate a break in immune tolerance and then accelerate the development of systemic and articular inflammation in susceptible individuals(2, 3). Studies of the preclinical phase of RA have demonstrated seropositivity for RA autoantibodies such as rheumatoid factor (RF) and anti-citrullinated protein antibodies (ACPA), elevations in pro-inflammatory cytokines and the presence of aberrant epigenetic processes (2–4).

A role for lifestyle habits and diet as risk factors for the development of RA has been proposed (5, 6). To date, the best documented lifestyle risk factor for RA development is tobacco smoking as described in multiple populations worldwide (7). Several dietary risk factors have also been studied in epidemiologic and cohort studies(8, 9). Since vitamin D deficiency has been shown to be prevalent in patients with early and established RA and an association with disease activity has been demonstrated, its potential role as a risk factor for RA development has been hypothesized (8, 10). Currently, there are limited studies that can define this association beyond any reasonable doubt. Existing literature suggests that lower vitamin D intake is associated with increased RA risk, while hypovitaminosis D affects disease activity score, bone-mass index, and regulatory T-cell proliferation in treatment-naïve RA patients(10).

Biologically synthesized inside the epidermis due a UV photolysis, vitamin D is hydroxylated by a family of cytochrome P450 enzymes called 25-hydroxylases to form 25(OH)D, the most predominant circulating form of vitamin D metabolite. 25(OH)D is further hydroxylated in the kidney and other tissues to form 1,25(OH)_2_D, the most biologically active form of vitamin D involved in maintenance of bone density, calcium absorption, and immunomodulation, including regulation of the metabolic phenotype of innate and adaptive immune cells (11). Vitamin D inhibits the differentiation of tissue-resident antigen-presenting cells (APCs) like dendritic cells (DCs) and macrophages from monocytes and prevents their maturation. Alternatively, it promotes the formation of tolerogenic DCs from bone marrow-derived DCs. Active vitamin D is also shown to affect the polarization and activation of CD4^+^ T lymphocytes, inhibit IL-17 production from Th17 cells. induced regulatory T-cells and inhibit the production of pro-inflammatory cytokines and chemokines. Additionally, 1,25(OH)_2_D also suppresses the differentiation of human B-cells and stimulates the production of immunomodulatory host-defense proteins such as LL-37 (8, 10, 12, 13)

In the current study, we evaluated the circulating levels of 25(OH)D, the active metabolite of vitamin D, in a cohort of first-degree relatives (FDR) of INA RA patients, and attempted to relate these levels to the onset of clinically detectable RA in individuals who ultimately developed disease. Our results suggest that, as with RA patients, ACPA seropositive FDR exhibit low levels of circulating 25(OH)D as a group, compared to ACPA seronegative FDR. Surprisingly, individuals who developed RA demonstrated an increase in 25(OH)D at the time of disease onset.

## METHODS

### Study Design

INA study participants were recruited from Cree, Ojibway, and Ojicree communities in Central Canada (14, 15). The Biomedical Research Ethics Board of the University of Manitoba approved the overall design of the study and consent forms (Ethics: 2005:093, Protocol: HS14453). The conduct of the study was guided by the principles of Community Based Participatory Research, a cornerstone of the Canadian Institutes of Health Research guidelines for Aboriginal health research (http://www.cihr-irsc.gc.ca/e/29134.html). The study participants provided informed consent after the study was explained to them in detail, with the help of an INA translator from their community, where necessary. For the cross-sectional study, the baseline serum samples of the following 3 groups were identified: (1) ACPA-positive RA patients, all of whom met the 2010 ACR/EULAR criteria, (2) ACPA-positive first-degree relatives (FDR) without any clinical evidence of joint or systemic inflammation and whose serum had detectable levels of ACPA (table 1), and (3) unaffected ACPA-negative FDR. Where possible, one sample from the winter and one from the summer for each FDR was included in the analysis. A subset of FDR who were followed longitudinally into RA onset, hereby referred to as “progressors”, had serum samples from several preclinical time points available for analysis, along with a sample from the study visit where they were first noted to have the onset of joint inflammation (referred as transition point). This was defined as the presence of one or more swollen joints deemed by the study rheumatologist (HEG) to represent active synovitis (16).

**Table 1:** Clinical characteristics of the study participants - All values are reported as median (range). RA = Rheumatoid Arthritis, RF = rheumatoid factor, anti-CCP = anti cyclic citrullinated protein antibody, CRP = C-reactive protein, NA = not applicable, BMI = body mass index.

|  | **ACPA-/FDR**<br>**(N=54)** | **ACPA+/FDR**<br>**(N=24)** | **RA Patients**<br>**(N=56)** | **Progressors**<br>**(N=14)** |
| --- | --- | --- | --- | --- |
| **Age, median** | 44 (23-7) | 46(25-64) | 47 (18-84) | 37 (27-76) |
| **Sex (female/male)** | 34/20 | 21/6 | 45/11 | 11/3 |
| **Age at transition,**<br>**median (range),**<br>**years** | - | - | - | 32 (23-68) |
| **Disease duration,**<br>**median (range)**<br>**years** | - | - | - | 5 (1-9) |
| **DAS28 scores,**<br>**median (range)** | NA | NA | 3.3 (1.4-5.8) |  |
| **CRP titer, mg/L,**<br>**median (range)** | 4.45 (1-16) | 2.3 (1-12) | 6 (1-85) | 7.2 (At onset) |
| **RF titer, IU/mL**<br>**median (range)** | - | 140(19-1870) | 327 (19-4110) | 294 (At Onset) |
| **Anti-CCP titer,**<br>**U/mL, median**<br>**(range)** | 3 (0-28) | 66(23-292) | 201 (19-330) | 300.5 (At Onset) |
| **BMI, kg/m<sup>2</sup>median**<br>**(range)** | 29 (21-54) | 28 (21-37) | 30.1(20-52) | 26.6 (At Onset) |

### Sample collection, storage and immunoassays

Venous blood was collected into SST™ serum separation tubes (BD Vacutainer Systems) and centrifuged for serum separation as per the manufacturer’s instructions. Serum was stored at −20°C until further use. C-reactive protein (CRP) levels were measured using a human high-sensitivity CRP (hs-CRP) ELISA kit (Biomatik, Canada) as per the manufacturer’s instructions. ACPA was detected using the BioPlex^®^ 2200 System anti-CCP reagent kit (Bio-Rad, US) and cutoff levels used were according to the manufacturer’s instructions. Concentrations of 25(OH)D (OKEH02569; Aviva Systems Biology) and VDBP were measured using ELISA kits as per the manufacturer’s instructions. For analysis, serum concentrations of 25(OH)D ≤ 75 nmol/L were considered to be vitamin D insufficient, while BMI < 30 were considered non-obese and CRP < 3mg/mL were considered normal (17).

### Data Analysis & Statistics

GraphPad Prism version 7.0 and SPSS 25.0 for windows (IBM Corp, USA) were used for data analysis and graphical data representation. Continuous variables were presented as mean ± SD. Non-parametric Kruskal-Wallis test with Dunn’s post-hoc method, repeated measures ANOVA with Greenhouse-Geisser correction, independent samples T-test or Wilcoxon matched pairs signed rank T-test were used for conducting comparisons as required and *P-*values < 0.05 were considered as statistically significant. Linear regression analysis was performed to identify an association model with predictive risk factors of RA.

## RESULTS

### Characteristics of the study participants

The study population consisted of RA patients (N=56), ACPA+/FDR (N=24), ACPA-/FDR (N=54), and FDR who progressed to develop RA while being followed in the study (progressors; N=14). The demographic and clinical characteristics of the study cohort are shown in Table 1. Most of the study cohort in all groups were females and the mean age was 47.8 ± 13.2years (mean + SD), although the mean age of the progressors was younger than the rest of the cohort. Not unexpectedly, RA patients, as a group, had higher CRP levels than the FDR groups. All of the RA patients were on disease-modifying anti-rheumatic drugs (DMARD) and the mean DAS28 score was 3.4 ± 1.2. The 14 progressors had a median age of 32 (23–68) years at the time they developed inflammatory arthritis.

### Effect of storage time and seasonal variation on circulating levels of 25(OH)D

As shown in Table 2a, mean serum concentrations of 25(OH)D were significantly higher in samples collected and stored after 2013 (storage period < 5yrs) compared to samples collected before 2013 (storage period > 5yrs). Based on this, we corrected for the duration of storage by excluding samples collected and stored before 2013.

**Table 2:** (a) Effect of storage time on the distribution of 25(OH)D levels between summer (April-September) and winter months (October – March). (b) Effect of season on the distribution of 25(OH)D levels in all study groups. SD = standard deviation

|  | Serum 25(OH)D concentrations in all study participants depending on season and storage time |  |  |  |  |  |  |  |  |  |  |  |
| --- | --- | --- | --- | --- | --- | --- | --- | --- | --- | --- | --- | --- |
|  | Summer |  |  |  |  |  | Winter |  |  |  |  |  |
|  | >=5y |  |  | <=5y |  |  | >=5y |  |  | <=5y |  |  |
|  | Total N | Mean | SD | Total N | Mean | SD | Total N | Mean | SD | Total N | Mean | SD |
| **Vit D (nmol/L)** | 30 | 19.20 | 16.12 | 101 | 100.66 | 57.36 | 31 | 16.84 | 11.97 | 25 | 107.49 | 53.49 |

(b)
|  | Serum (OH)D concentrations in summer and winter months in all study participants |  |  |  |  |  |
| --- | --- | --- | --- | --- | --- | --- |
|  | Summer |  |  | Winter |  |  |
|  | Total N | Mean | SD | Total N | Mean | SD |
| **Vit D (nmol/L)** | 131 | 82.00 | 61.40 | 55 | 57.85 | 58.65 |

When we analyzed only samples obtained after 2013, we further observed a trend towards higher mean 25(OH)D levels during summer months (April-September) compared to winter months (October-March) in the ACPA-FDR group (140.29 +/− 78.18 vs 110.8 +/− 51.91 (mean +/− SD); *P*=0.129). Based on these considerations, our subsequent comparisons between groups were based exclusively on samples gathered after 2013 during the summer months.

### Circulating levels of 25(OH)D are lower in RA and ACPA+ FDR compared to ACPA-FDR

We compared circulating 25(OH)D levels in RA patients, ACPA+ FDR, and ACPA-FDR (figure 1). As a group, RA patients and ACPA+ FDR demonstrated significantly lower levels compared to the ACPA-FDR group (87.48 +/− 32.76 vs 87.98 +/− 59.06 vs 125.86 +/− 67.59 (mean +/− SD); *P* = 0.001). There was no significant difference between RA patients and ACPA+ FDR in their circulating levels of 25(OH)D.

**Figure 1.**
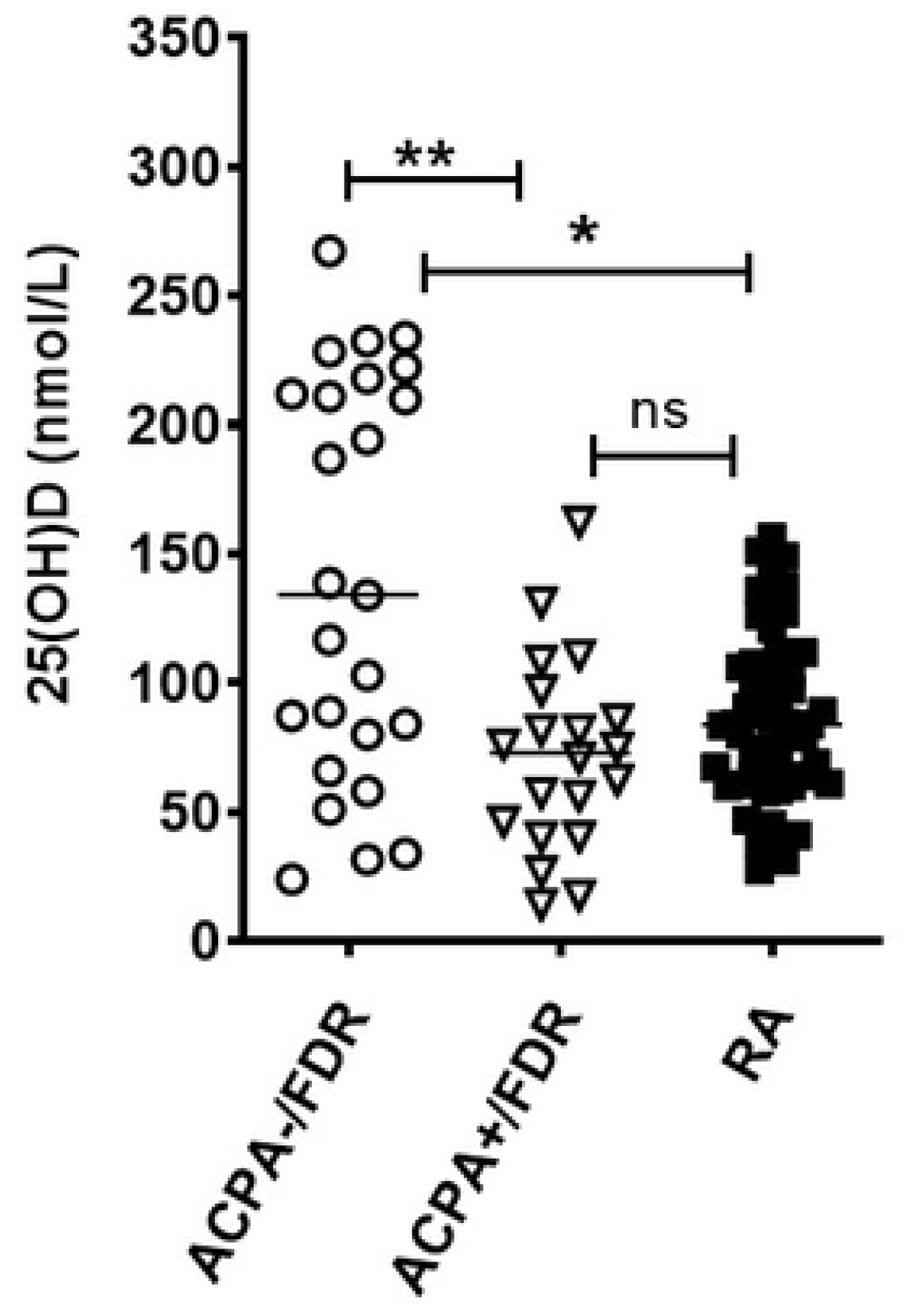
Cross-sectional analysis of 25(OH)D levels in ACPA-/FDR, ACPA+/FDR and RA patients. Scatter plot showing the distribution of 25(OH)D levels after correcting for storage and seasonal effect. Data was analyzed by Kruskal-Wallis test with Dunn’s post-hoc test (\*\**P*<0.01, \**P*<0.05)

We performed a linear regression analysis to explore potential associations between circulating levels of 25(OH)D and preclinical RA risk factors. In this analysis, circulating 25(OH)D had a significant inverse association with anti-CCP antibody levels (β = −0.317; *P* = 0.005) and positive association with RF antibody levels (β = 0.266; *P* = 0.017). No other association reached statistical significance (table 3).

**Table 3:**
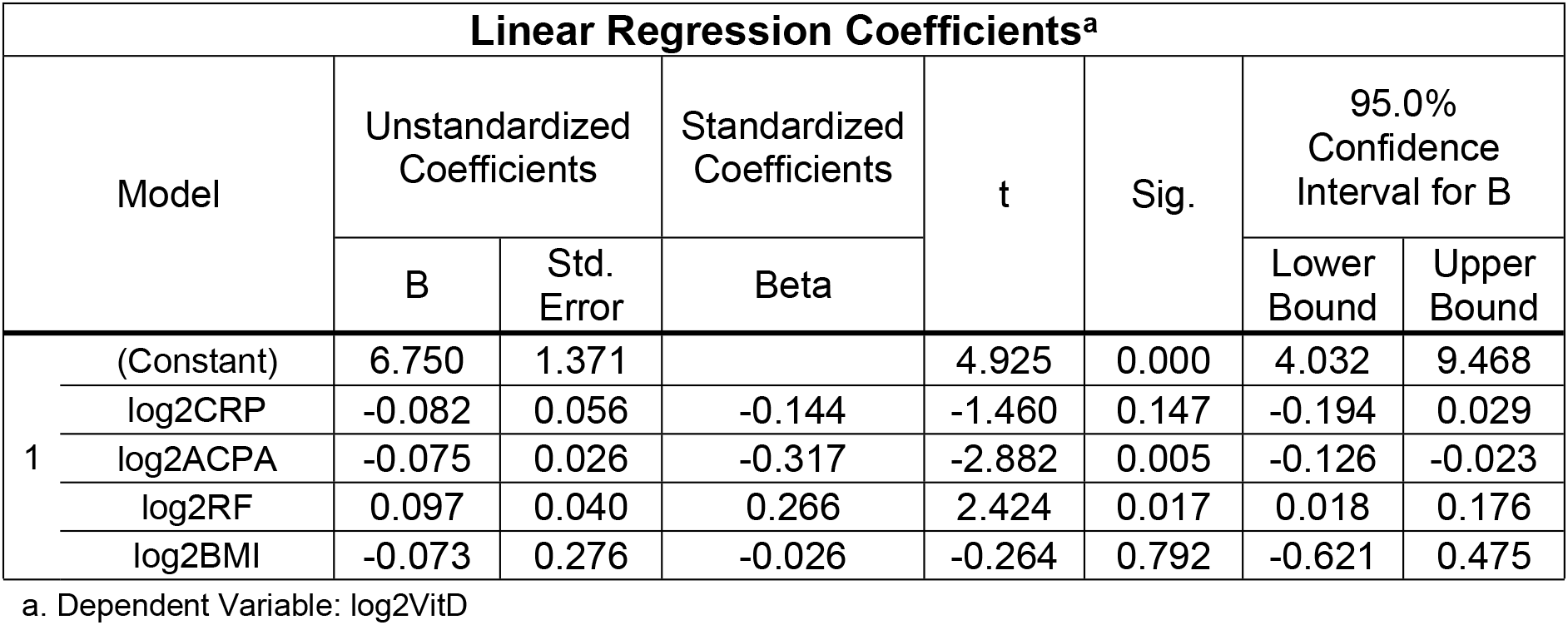
Linear regression analysis to demonstrate the relationship between serum 25(OH)D levels (dependent variable) and clinical risk factors associated with RA (independent variables). Analysis was performed on log_2_ values.

### Circulating levels of 25(OH)D rise in FDR who progressed to develop RA

We tested all available longitudinal samples (before and after 2013) from 14 progressors who ultimately developed RA for levels of 25(OH)D and VDBP (figure 2). Samples obtained before (labelled as T-5, T-4, T-3, T-2 and T-1), and the sample obtained at the time of IA diagnosis (called as T0) were tested for each progressor. Compared to both pre-RA samples (T2 and T1), 25(OH)D levels in the RA onset sample (T0) increased over time in most progressors (*P* = 0.001) as they approached IA transition point (fig 2A and 2B). 25 (OH) D were significantly higher at T0, compared to pre-transition points T-1 and T-2 (*p* = 0.0010 and *p* = 0.0017 respectively). Mean time difference between points −1 and 0 was ~19.79 + 10.16 months (mean + SD), while the time difference between points −2 and 0 was ~44.75 + 20.18 months (mean + SD) respectively (fig 2B).

**Figure 2.**
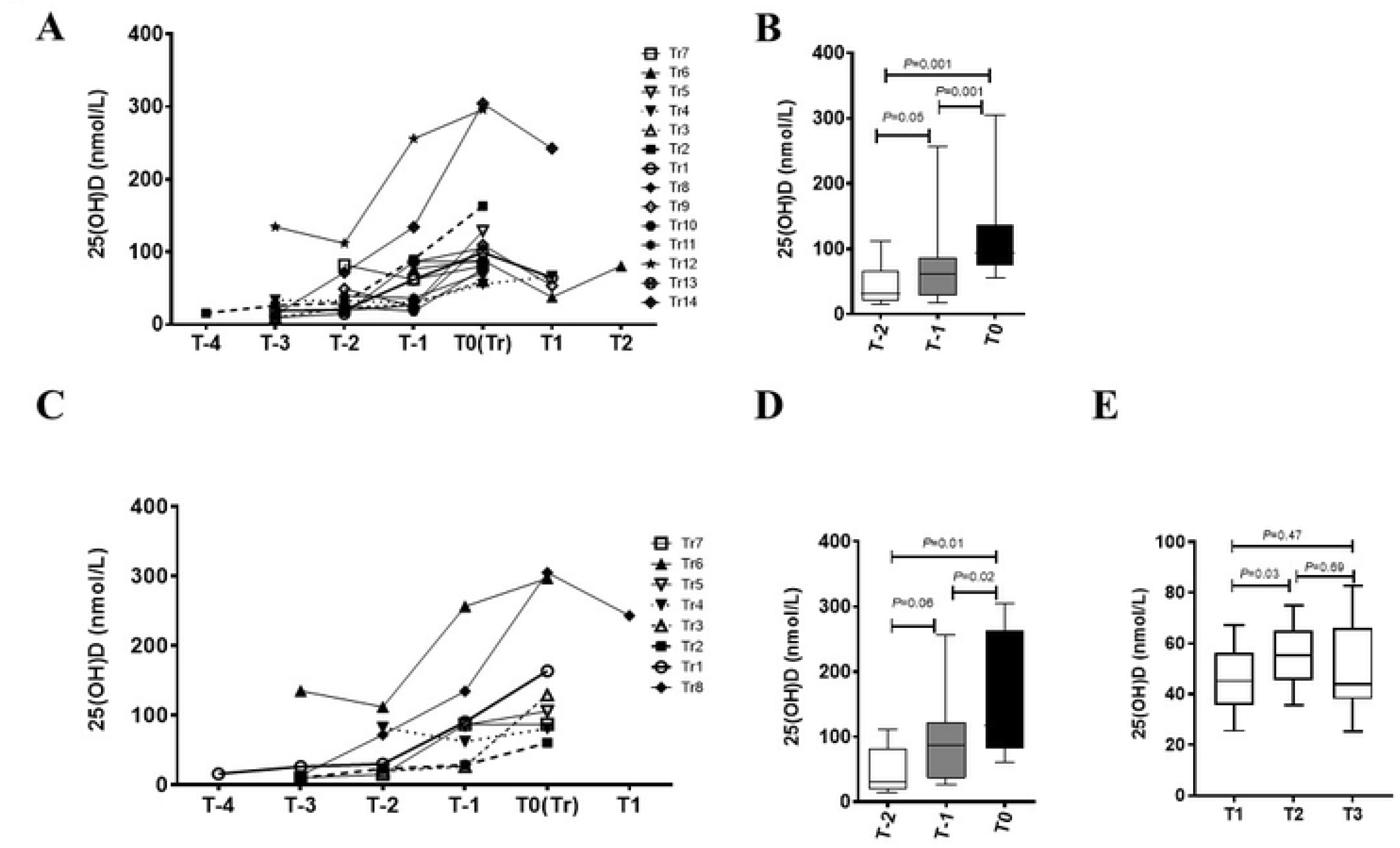
Longitudinal analysis of 25(OH)D levels in progressors. (A) Line graph showing the evolution of 25(OH)D levels over time in progressors (N=14) prior to RA onset (T-1 to T-4), at the time of clinical diagnosis of RA onset (T0) and post-onset (T1 and T2) (B) Box-Whiskers plot showing the distribution of 25(OH)D at two time points prior to RA onset (T-1 and T-2) and at transition point (0). Data was analyzed by analyzed by repeated measures ANOVA using Greenhouse-Geisser model (C) After correcting for storage effect, line graph showing the evolution of 25(OH)D levels over-time in progressors (N=8) prior to RA onset (T-1 to T-4), at the time of clinical diagnosis of RA onset (T0) and post-onset (T1 and T2). (D) Box-Whiskers plot showing the distribution of 25(OH)D at two time points prior to RA onset (T-1 and T-2) and at transition point (T0), in samples collected after 2013. (E) Box-Whiskers plot showing the distribution of 25(OH)D in ACPA-/FDR collected at 3 different time points. Data was analyzed by repeated measures ANOVA with Geisser-Greenhouse correction.

Because of the duration of storage effect described above, we further analyzed only samples from progressors who had all of their samples collected after 2013 (n=8). In this subset, the mean time difference between T-1 and T0 was ~18.75 + 12.19 months (mean + SD), while the time difference between T-2 and T0 was ~50.0 + 25.14 months (mean + SD) and 29.71 + 16.3 months (mean + SD) between pre-transition points −1 and −2 respectively. This analysis showed that the rise in 25(OH)D levels at the time of IA diagnosis was clearly evident in this more recently studied subset (fig 2C and 2D; *p* = 0.01 and *p* = 0.02). To further confirm that this rise in 25(OH)D levels at the time of IA onset was specific, we analyzed longitudinal samples from a subset of ACPA-FDR collected at multiple intervals (fig 2E). These results indicate that that 25(OH)D concentrations in these individuals remained relatively stable over time indicating that the phenomenon observed in the progressor is specific to that subset.

As the bioavailability and biological activity of 25(OH)D is dependent on the circulating levels of VDBP with 85% of 25(OH)D being bound to VDBP(18), we tested VDBP levels in the same samples to determine whether the levels of this glycoprotein followed a similar pattern to the free 25(OH)D that were measured. This analysis showed that VDBP levels remained stable over time (fig 3A and 3B; *p* = 0.432), even as 25(OH)D increased in the progressor group.

**Figure 3.**
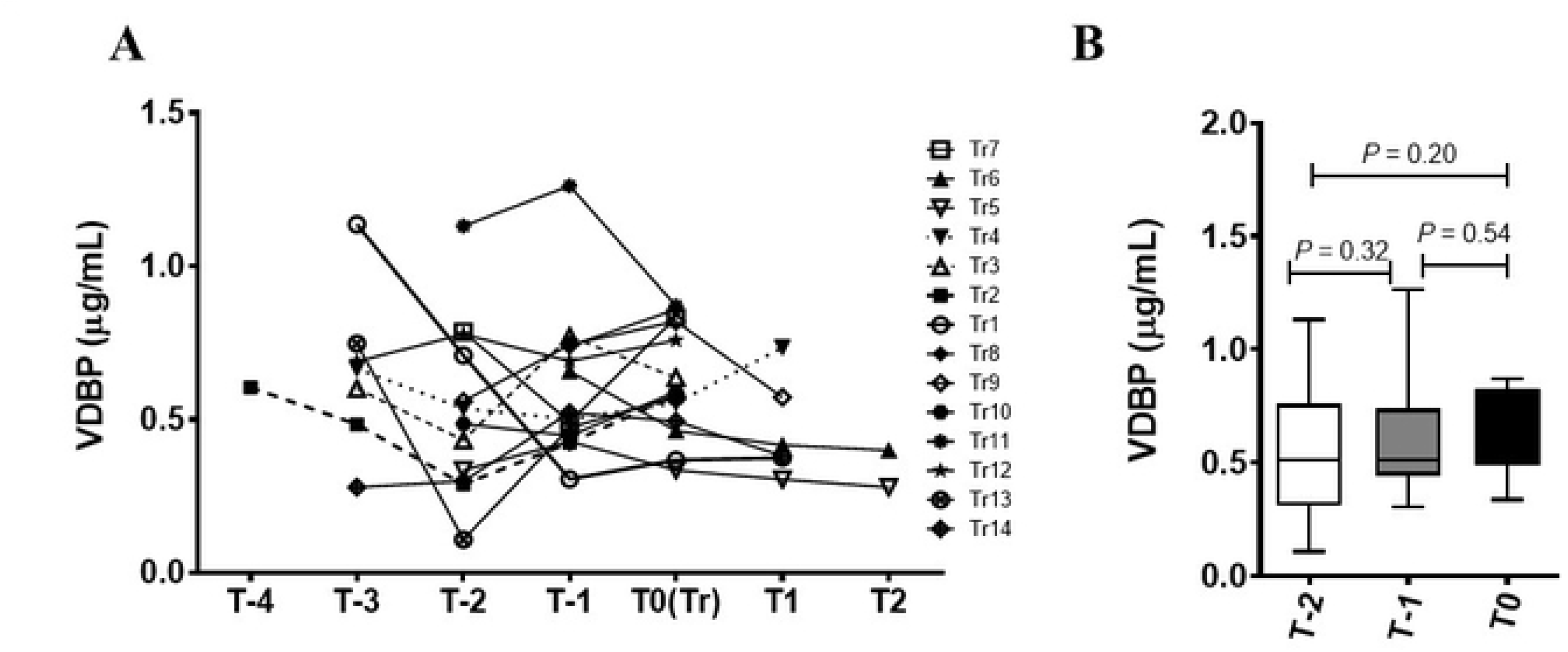
Longitudinal and cross-sectional analysis of VDBP. (A) Line graph showing the evolution of VDBP levels over-time in progressors (N=14) prior to RA onset (T-1 to T-4), at the time of clinical diagnosis of RA onset (T0) and post-onset (T1 and T2) (B) Box-Whiskers plot showing the distribution of VDBP at two time points prior to RA onset (T-1 and T-2) and at transition point (T0). Data was analyzed by analyzed by repeated measures ANOVA using Greenhouse-Geisser model.

## DISCUSSION

We surveyed the serum levels of 25(OH)D in a cohort of INA RA patients and their unaffected FDR having no clinically detectable joint inflammation. We tested 25(OH)D levels at different preclinical stages of disease evolution, both cross-sectionally and longitudinally. The cross-sectional analysis provided insights into the relationship between 25(OH)D levels and RA autoantibodies, in the presence and absence of clinical disease. A longitudinal analysis of preclinical samples from individuals who eventually developed IA provided the unique opportunity to examine the association of vitamin D status and disease onset.

We quantified free 25(OH)D levels in circulation as an indicator of vitamin D status. Considered as the predominant circulating form of vitamin D, free/unbound 25(OH)D concentrations in serum are directly reflective of the available vitamin D while having a longer half-life compared to other vitamin D metabolites, making it well suited to assess the overall vitamin D status (19). While ~0.01-0.03% free unbound 25(OH)D is available in the circulation, 85-90% 25(OH)D is bound to VDBP and 10-15% exists in an albumin-bound form (11). Circulating levels of free 25(OH)D are known to fluctuate based on several factors such as season, dietary intake of vitamin D, and intensity of ultraviolet radiation in sunlight (need citations). While we were unable to account for all of these variables in our analysis, a number of them were particularly relevant to the findings of this study.

Unexpectedly, we demonstrate that in a cohort of longitudinally followed individuals who ultimately developed IA, 25(OH)D levels consistently increased as they approached disease onset, with the highest concentrations being detected at the time they were deemed to have the onset of IA. This finding was demonstrated in almost all of the “progressors” who were studied. The 25(OH)D levels did not decrease in any of these individuals, although the magnitude of the increase varied considerably. Moreover, there was no consistent change in the levels of the carrier protein VDBP in the longitudinal sample analysis. This intriguing increase in 25(OH)D levels at disease onset is particularly noteworthy since our cross-sectional analysis of patients with established RA demonstrated significantly lower levels of 25(OH)D compared to a cohort of seronegative, unaffected controls. The latter finding echoes the results of other cross-sectional studies of RA patients and controls (20–24).

Circulating 25(OH)D concentrations in RA patients are known to be associated with the presence of ACPA and RF (25, 26). However, current evidence is conflicting on the role of 25(OH)D and vitamin D during the preclinical stages of RA (27, 28). In our INA cohort, we show that 25(OH)D concentrations in seropositive FDR, as with RA patients, are lower than in seronegative FDR. Longitudinal analysis of 25(OH)D levels in progressors demonstrates that there is a consistent increase in levels at the onset of inflammatory arthritis compared with earlier time points.

In attempting to explain this unexpected finding, several possibilities can be considered. One important consideration is the potential confounding effects of duration of sample storage and seasonal variation on 25(OH)D levels. It is conceivable that 25(OH)D levels “appeared” to increase because of a decay in this analyte in older stored samples from the same individuals. We clearly demonstrate this duration of storage effect in samples stored > 5 years, with older samples having lower levels. Our data contrasts that of previous publications suggesting that duration of sample storage had minimal impact on 25(OH)D stability (29, 30), although this decay may be less likely to occur if samples are stored at −80°C. Our analysis of the subgroup of 8 progressors whose samples were all gathered within the most recent five-year timeframe all showed increasing 25(OH)D levels at the time of IA onset. Moreover, this increase in levels was not observed in the longitudinal samples of seronegative individuals who did not develop IA. Together, these observations suggest that this phenomenon cannot be explained on the basis of time in storage of the samples.

We also demonstrate that samples gathered during the winter months, on average, exhibit lower levels of 25(OH)D compared to samples gathered in the summer months, results that are consistent with published literature in other populations, including in other INA communities (31, 32). Of note, we show that a majority of our study participants had summer concentrations of 25(OH)D that were greater than 75nmol/L, with only 34% demonstrating levels deemed to be in the vitamin D insufficiency range. These results are consistent with studies in other INA communities (33). Nevertheless, our analysis suggests that seasonal variation does not account for the observed rise at the time of IA onset. Factors like increased exposure to sunlight or dietary intake of vitamin D supplements are insufficient as 25(OH)D levels remained consistently increased over a long period of time extending up to ~19months preceding onset of clinically detectable symptoms.

Previously, we showed that single-nucleotide polymorphism Fok1 (rs2228570) within the VDR gene is significantly associated with RA onset in INA population (34). Additionally, systemic inflammation, obesity, and parathyroid hormone levels have been associated with changes in circulating vitamin D levels. (12, 35–40). These factors can regulate the concentrations and/or the activity of 25-D-hydoxylases and 1,25-D-hydroxylases involved in vitamin D metabolism leading to either an impaired cellular uptake of 25(OH)D from circulation or conversion to metabolically active 1,25(OH)_2_-D. Additional investigations are warranted to identify how these factors might regulate the mechanisms underlying increased 25(OH)D concentrations.

In summary, we show that 25(OH)D levels are reduced in seropositive INA RA patients and unaffected seropositive FDR. Surprisingly we demonstrate a consistent increase in 25(OH)D levels at the time of onset of inflammatory arthritis compared to earlier preclinical time points. Future studies are required to understand the basis for this increase in 25(OH)D levels and the potential link with the break in immune tolerance preceding RA onset.

## ACKNOWLEDGEMENTS

We acknowledge the contribution of study participants from indigenous communities who donated blood for our study. We also recognize Chief and Band Councils of Norway House and St. Theresa Point Manitoba for their invaluable cooperation.

